# Modelling fibrillogenesis of collagen-mimetic molecules

**DOI:** 10.1101/2020.06.08.140061

**Authors:** A. E. Hafner, N. G. Gyori, C. A. Bench, L. K. Davis, A. Šarić

**Affiliations:** University College London; Univ. College of London, UK

## Abstract

One of the most robust examples of self-assembly in living organisms is the formation of collagen architectures. Collagen type I molecules are a crucial component of the extracellular-matrix where they self-assemble into fibrils of well defined striped patterns. This striped fibrilar pattern is preserved across the animal kingdom and is important for the determination of cell phenotype, cell adhesion, and tissue regulation and signalling. The understanding of the physical processes that determine such a robust morphology of self-assembled collagen fibrils is currently almost completely missing. Here we develop a minimal coarse-grained computational model to identify the physical principles of the assembly of collagen-mimetic molecules. We find that screened electrostatic interactions can drive the formation of collagen-like filaments of well-defined striped morphologies. The fibril pattern is determined solely by the distribution of charges on the molecule and is robust to the changes in protein concentration, monomer rigidity, and environmental conditions. We show that the fibril pattern cannot be easily predicted from the interactions between two monomers, but is an emergent result of multi-body interactions. Our results can help address collagen remodelling in diseases and ageing, and guide the design of collagen scaffolds for biotechnological applications.

**Statement of Significance:** Collagen type I protein is the most abundant protein in mammals. It is a crucial component of the extracellular-matrix where it robustly self-assembles into fibrils of specific striped architectures that are crucial for the correct collagen function. The molecular features that determine such robust fibril architectures are currently not well understood. Here we develop a minimal coarse-grained model to connect the design of collagen-like molecules to the architecture of the resulting self-assembled fibrils. We find that the pattern of charged residues on the surface of molecules can drive the formation of collagen-like fibrils and fully control their architectures. Our findings can help understand changes in collagen architectures observed in diseases and guide the design of synthetic collagen scaffolds.

---

Reversible protein assembly into fibrils and networks is crucial for biological function: protein filaments give shape and support to cells, remodel cells, and bind them into tissues. One of the most remarkable examples of fibrillar assembly in nature is the assembly of collagen proteins, the most abundant proteins in animals [1]. Collagen molecules are secreted by cells as monomers, followed by their spontaneous organisation into fibrils, bundles, and networks that span from molecular to macroscopic length-scales. Collagen molecules are the structural basis of mammalian connective tissues, including those of the heart, skin, and bones. Beyond its structural role, collagen is also the major component of the extracellular matrix – the dynamical network of fibrils that surrounds cells – where it plays a key role in the determination of cell phenotype, cell adhesion, tissue regulation and signalling [2, 3].

It remains unclear which properties of collagen molecules drive their assembly into robust hierarchical structures. Here we design a minimal coarse-grained computer model that enables us to explore the physical and chemical principles of collagen assembly, hereby bridging molecular and mesoscopic scales.

Each collagen monomer is about 300nm long and 1.5nm in diameter, and consists of three peptide chains spiralling around each other in a form of triple helix [1]. In the case of type I, the most abundant collagen type, such triple-helical monomers longitudinally stagger into fibrils that display a regular array of gap and overlap areas (Figure 1a). This periodicity of collagen type I fibrils, called the D-spacing or D-periodicity, has a characteristic length of ∼67 nm, and is remarkably conserved across the animal kingdom [4]. Fibrils then associate via specific cross-links into fibres which can be up to 10 *µ*m-thick and few mm long, and interconnect into networks. Fibrils very much alike to those formed in live cells are regularly reconstituted in-situ, without the presence of other factors normally present in cells. Therefore, the formation of collagen fibrils is believed to be a robust self-assembly process [1, 2].

**FIG. 1:**
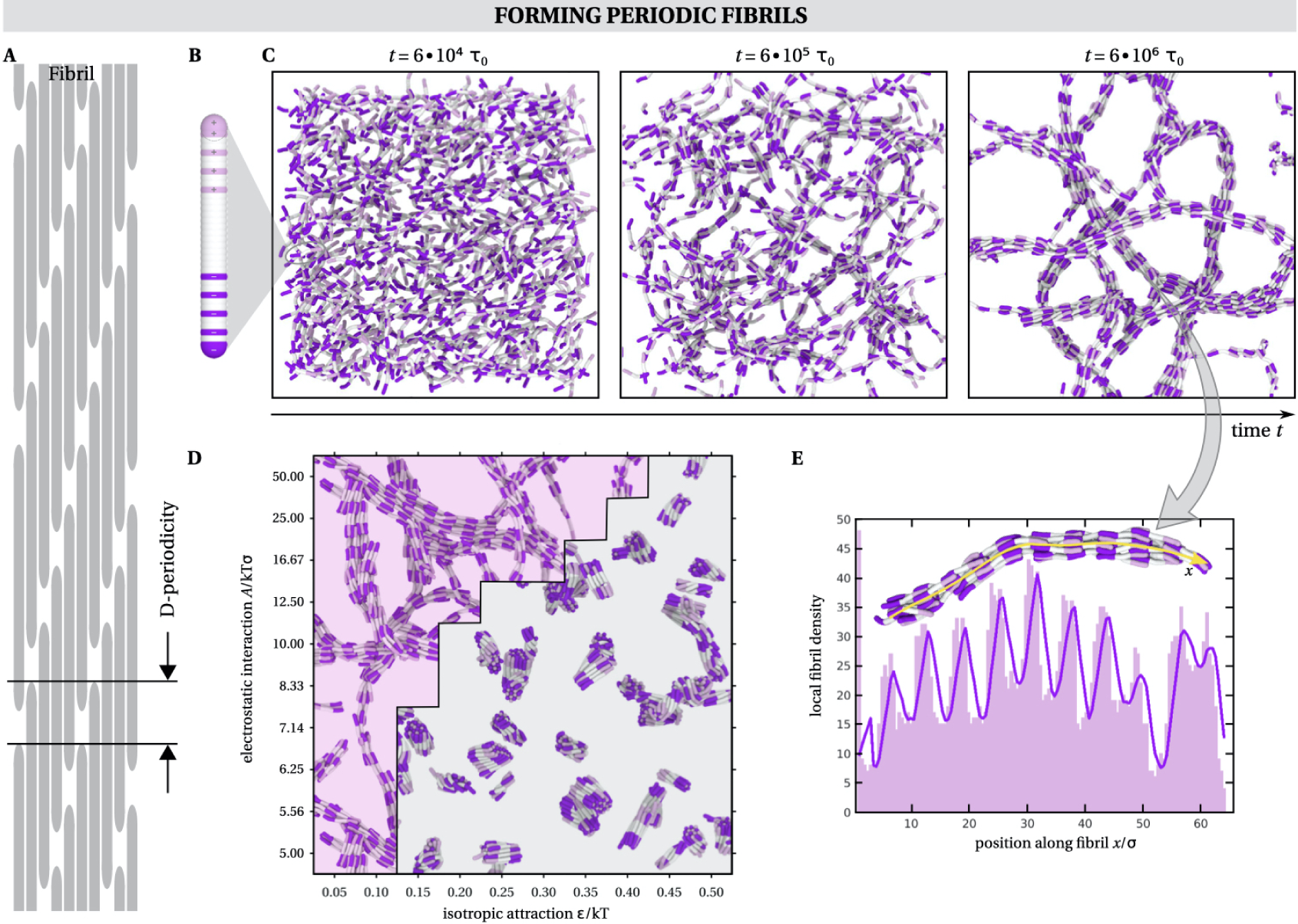
Modelling collagen fibrillogenesis. **a)** Collagen type I molecules self-assemble into fibrils of a characteristic pattern with gap and overlap regions, termed D-periodicity. **b)** A single collagen-mimetic molecule simulated here consists of 36 beads with a charge distribution as indicated, resembling the one of D-mimetic synthetic peptide [7]. **c)** Simulation snapshots showing the dynamic self-assembly of such molecules into periodic fibrils for isotropic attraction *epsilon* = 0.05*kT*, electrostatic interaction *A* = 10*kT σ*, monomer rigidity *k*_*angle*_ = 50*kT*, and molecule concentration *c*_*mol*_ = 0.001*σ*^−3^. **d)** Phase diagram of fibrillogenesis as a function of the strength of electrostatic interactions *A* and the isotropic attraction *ϵ*. Periodic fibrils are only formed when the electrostatic interactions dominate, while otherwise isolated clusters are formed. **e)** In order to analyse the periodicity of the fibrils, the local density along the fibril backbone (indicated with a yellow line) is measured. The periodicity is defined as the mean distance between peaks of the density profile.

To probe the principles of the assembly, smaller synthetic peptides, which mimic the structure and periodicity of collagen molecules, have been developed [5–9]. The arrangement of charged residues has been suggested to be crucial for the correct arrangement of collagen-like molecules [1, 10]. For synthetic collagen-mimetic peptides, the placement of charged amino-acids along the chain of the molecule has been shown to influence the nature and morphologies of the resulting assemblies, which include crystals, fibrils, and networks [5– 9]. Certain arrangements of charges were found to give rise to fibrils with pronounced offsets [1, 7] that qualitatively resemble the D-spacing of native collagen. In the case of native type I collagen, about 15-20% of all residues are charged at physiological pH, and the fibril morphologies have been found to strongly depend on the ionic strength of the solution [6, 11–13, 13]. The morphologies also dramatically change upon fibril glycation, a process in which sugar molecules bind to fibrils and neutralise native surface charges [14]. In addition to the importance of charges, it has been shown that fibrillogenesis of native collagen is promoted by warming [15], pointing to the importance of hydrophobic interactions in driving the assembly [1, 10]. However, the quantitative relationship between the charge arrangement, hydrophobicity, and the resulting self-assembled phase is not known, nor is the influence of the charge patterns on the presence and the value of the fibril periodicity, and fibril sizes and morphologies.

From the modelling point of view, atomistic and coarse-grained models have been used to study the stability and dynamics of single collagen molecules, pre-assembled collagen fibrils and crystals [16, 17], as well as their mechanical properties [18–22]. However the dynamic assembly of collagen and collagen-mimetic molecules from solution has not been yet addressed due to the large system sizes required for such simulations.

Here we developed a minimal model for the assembly of collagen-mimetic molecules. In the model the molecules are described as elastic rods decorated with a pattern of charges. Two such rods can interact via screened electrostatic interactions as well as via generic, hydrophobic, attractions. We run molecular dynamics (MD) simulations and devise a simple analytical model to establish the connection between the molecular design and the architecture of the resulting assemblies. We find that the interplay between electrostatic interactions and non-specific attractions can drive the system to the formation of dispersed clusters, or indeed, long interconnected fibrils of well defined D-periodicity. We show that the periodicity is extremely robust and is determined by the pattern of the charged residues along the monomer backbone. We show that the fibril morphology cannot be easily predicted from the properties of a single monomer or from the interactions between two monomers only. Using a simple analytical model, we demonstrate that the emergent fibril pattern can be well predicted from the interactions of a single monomer with the field created by its neighbouring molecules.

## Molecular dynamics model

In order to investigate the fibrillogenesis of collagen-mimetic molecules, we first develop a minimal coarse-grained model and perform molecular dynamics simulations. We base our initial model on the synthetic collagen-mimetic molecule, termed “D-mimetic”, which was shown to self-assemble into fibrils of defined D-periodicity [7]. Since this D-mimetic peptide consists of 36 amino acids, for the ease of mapping, our molecules consists of 36 beads connected into a linear chain. Each bead has a diameter *r*=1.12*σ*, where *σ* is the MD unit of length, and is connected to its neighbour by a harmonic bond *E*=*κ*_bond_(*r* − *r*_0_)^2^ with bond strength *κ*_bond_=500*kT* and equilibrium distance *r*_0_=0.255*σ*, resulting in a molecule length *l*=10*σ*. Consequently 1*σ*=1nm because the D-mimetic molecule has a length of 10nm. The rigidity of the molecule *κ*_angle_ is defined by an angular potential that acts between three adjacent beads, *E*=*κ*_angle_(*θ* − *θ*_0_)^2^ with *θ*_0_=*π*. We explore the influence of molecular rigidity *κ*_angle_ on the fibril structure. Furthermore, each bead carries a unit charge in accordance to the charge distribution of the D-mimetic peptide, as indicated in Figure 1 b.

The beads on different molecules interact via a generic attractive potential, described via a cut-and-shifted Lennard-Jones potential *E*_*LJ*_ = 4*ϵ*[(*σ/r*)^12^ − (*σ/r*)^6^] if the two beads are at distances shorter than *r* < *r*_*c*_ = 2*σ*. In this framework *ϵ* represents the strength of non-specific attractions and will be one of the parameters we will explore.

On top of this potential, two beads *i, j* that carry charge interact via a cut-and-shifted screened electrostatic potential (DLVO) 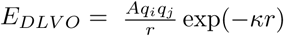 if the distance between them is shorter than *r* < *r*_*c*_ = 2*σ*. The screening length *κ* was chosen to be 1*σ*, which is equivalent to ∼1nm. This corresponds to the Debye screening length at physiological conditions. *q*_*i*_ carries only the information on the sign of the charge of bead *i* (±1), while the prefactor *A* determines the effective strength of the electrostatic interactions and it is another parameter we explore. To prevent nonphysically large forces due to the proximity of the like charges in a molecule and the overlaps in the volume of the neighbouring beads, we have excluded bead-bead interactions between 1-2, 1-3, 1-4, and 1-5 neighbours in the same molecule.

To run molecular dynamics simulations we place an ensemble of *N* = 1000 individual molecules into a 3D box of boxlength *L*, which results into a molecule concentration *c*_mol_=*N/L*^3^. The system was integrated in the *NV E* ensemble (*V* being the volume of the box and *E* the total energy of the system) with Langevin thermostat to capture Brownian motion of the molecules, within the LAMMPS molecular dynamics package [23]. The integration timestep was chosen to be 0.002*τ*_0_, *τ*_0_ being the MD unit of time, and the damping coefficient was set to 1 *τ*_0_.

## Simulating fibrillogenesis

We first set out to explore the assembly of “D-mimetic” collagen-mimetic monomers, which carry equally sized regions of positive and negative charge on two of its ends, separated by an equally long neutral area in its middle (Figure 1b). Such synthetic monomers have been found to create fibrils that display D-periodicity reminiscent of that of natural collagen fibrils [7]. We vary the strength of non-specific hydrophobic interactions that act along the length of the molecule, represented by *ϵ*, and the strength of screened electrostatic interactions, represented by *A* (while keeping the range constant).

As shown in Figure 1d, we find that when the isotropic interactions dominate, separated clusters of molecules are formed, which can possess a certain degree of crystalline packing, but do not display elongated morphologies. However, when the screened electrostatic interactions dominate, fibrils of highly periodic structures readily form, as shown in Figure 1c and d. The crossover between the two phases is rather continuous and mixed phases can occur around the border.

Interestingly, in the region of the phase space where the fibrils do assemble, they all appear to have the same morphology, characterised by periodic regions of gaps and overlaps. To analyse this periodicity of the self-assembled fibrils we measure the fibril mass density along the fibril backbone. As shown in Figure 1e for an example of one fibril, the density shows a well-defined periodic pattern of regions of high density (overlap) and regions of low density (gap). The periodicity *p* is then defined as the mean distance between the peaks of the density profile.

Figure 2a shows that the periodicity of the fibrils assembled for a certain combination of infractions is typically Gaussian distributed. Remarkably, we find that this mean value of the periodicity does not change substantially, neither in time (Figure 2b), nor as a function of the isotropic attraction *ϵ* (Figure 2c), the strength of screened electrostatic interactions *ϵ* (Figure 2d), the molecule rigidity *κ*_angle_ (Figure 2e), or the molecule concentration *c*_mol_ (Figure 2f). This indicates that the fibril periodicity is a robust feature, encoded purely in the molecular design.

**FIG. 2:**
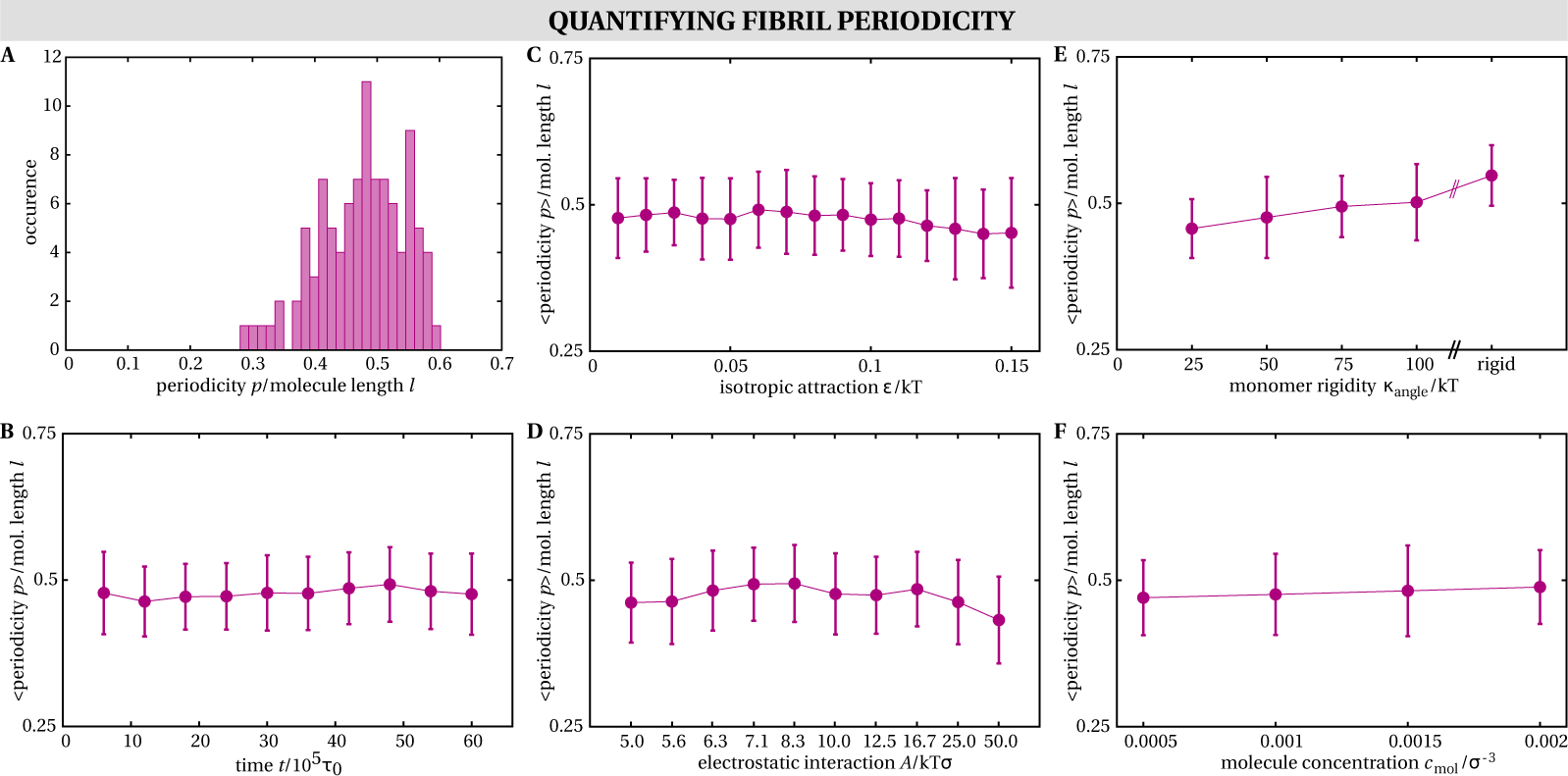
Analysing fibril periodicity. **a)** Distribution of periodicity *p* scaled by the molecule length *l* for fibrils that are formed by isotropic attraction *ϵ*=0.05*kT*, electrostatic interaction *A* = 10*kT σ*, monomer rigidity *κ*_angle_=50*kT*, and molecule concentration *c*_mol_=0.001*σ*^−3^. **b)** Scaled fibril periodicity as a function of time *t* for *ϵ*=0.05*kT*, interaction *A* = 10*kT σ, κ*_angle_=50*kT*, and *c*_mol_=0.001*σ*^−3^. **c)** Scaled Fibril periodicity as a function of the isotropic attraction *ϵ* for *A*=10*kT σ, κ*_angle_=50*kT*, and *c*_mol_=0.001*σ*^−3^. **d)** Scaled fibril periodicity as a function of the strength of the electrostatic interaction *A* for *ϵ*=0.05*kT, κ*_angle_=50*kT*, and *c*_mol_=0.001*σ*^−3^. **e)** Scaled fibril periodicity as a function of the monomer rigidity *κ*_angle_ for *ϵ*=0.05*kT, q*=10*kT σ*, and *c*_mol_=0.001*σ*^−3^. **f)** Scaled fibril periodicity as a function of the molecule concentration *c*_mol_ for *ϵ*=0.05*kT, A*=10*kT σ*, and *κ*_angle_=50*kT*.

## Predicting fibril periodicity

By the minimal design of our model, the only information that our molecules carry is stored in the pattern of charges on its surface. Hence the robust fibril periodicity in our model must emerge from the arrangement of charges on the monomer surface. We then ask the question: given the pattern on charges on the molecular surface, can we predict the value of the resulting fibril periodicity out of simple arguments?

Since the fibril periodicity appears not to depend on fine details of the monomer, we simplified our model and investigated the energy of a minimal, two-stranded fibril that is composed of a repetitive continuation of two molecules, as shown in Figure 3a. The fibril geometry and its energy are defined by three parameters: the tip-to-tip distance *radial gap* between two molecules in the same row, the surface-to-surface distance *lateral gap* between two molecules in neighbouring rows, and the *offset* between two molecules in the neighbouring rows. These molecules are rigid, but ineract via the same Lennard-Jones and Coulomb-Debye potentials as in our MD model. For each set of parameters – the strength of isotropic attraction *ϵ* and screened electrostatic interaction *A* – we minimise the energy of a single focus molecule in the field of the other molecules as a function of *radial gap, offset* and *lateral gap*.

**FIG. 3:**
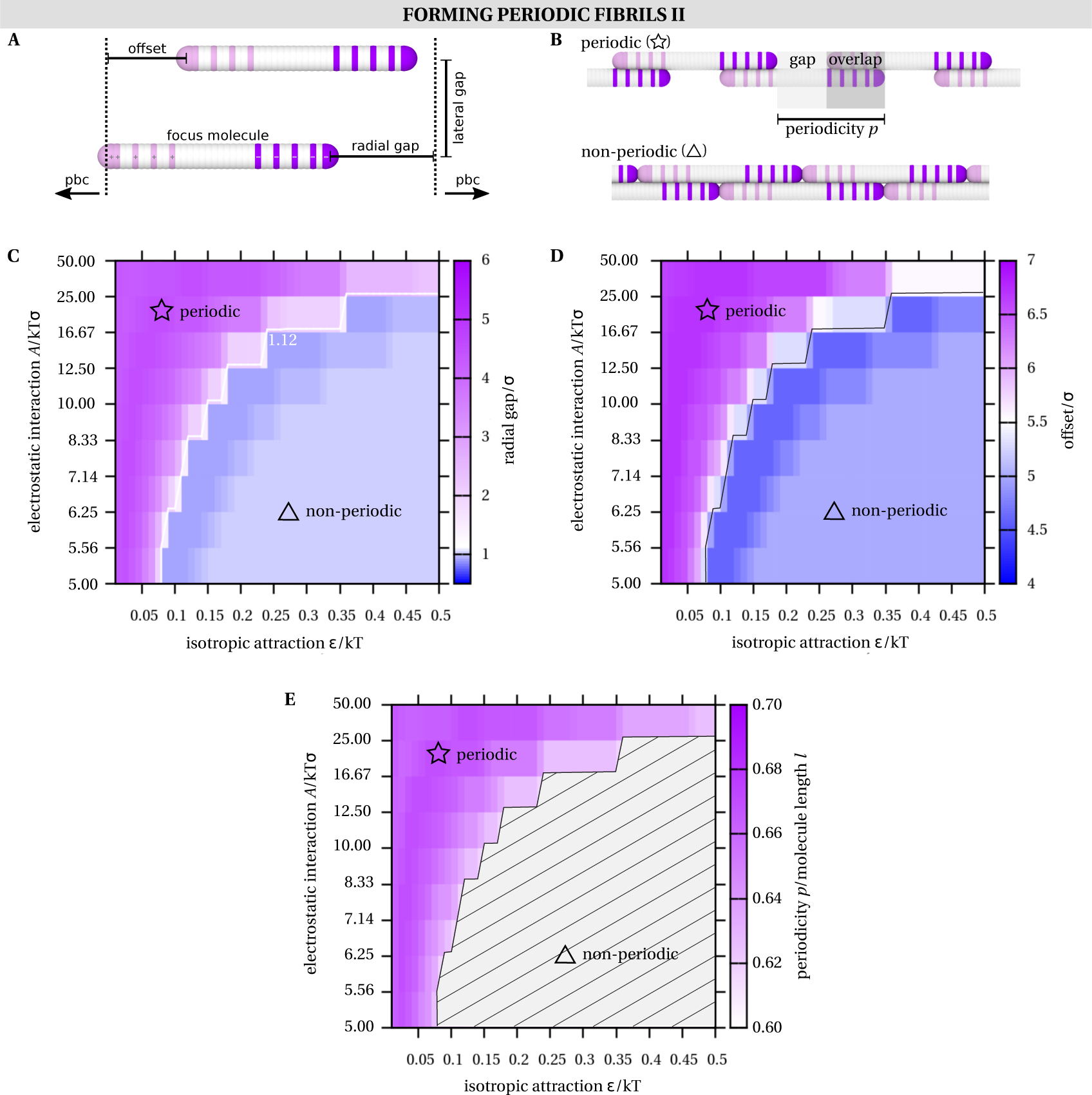
Analytic model for determining fibril formation and fibril morphologies. **a)** Schematic figure of the analytic model developed to predict the impact of the single molecule charge distribution on the fibril periodicity. For that purpose, the energy of the focus molecule (in the field that is created by its surrounding molecules found in a minimal two-stranded fibril) is measured and minimised as a function of the three geometrical parameters *offset, radial gap*, and *lateral gap*. **b)** For the charge distribution of the D-mimetic molecule either D-periodic or non-D-periodic configurations are found. The periodicity is defined as the sum of one gap and one overlap region. **c)** Value of *radial gap* that minimises the energy of the system as a function of the strength of the electrostatic interactions *A* and isotropic attraction *ϵ*. The *radial gap* determines if a fibril is D-periodic (*radial gap*> 1.12*σ*) or not (*radial gap*< 1.12*σ*). **d)** Value of *offset* that minimises the energy of the focus molecule as a function of electrostatic interaction *A* and isotropic attraction *ϵ*. **e** Periodicity of D-mimetic fibrils predicted by the analytic model as a function of the electrostatic interactions *A* and isotropic attraction *ϵ*. Remarkably, the periodicity does not substantially change throughout the phase space.

We find that the energy is minimised when *lateral gap*= 1.12*σ*, corresponding to the contact distance between the beads, for all the designs, hence we do not explore this geometric parameter further. In Figure 3c and 3d we show the values of radial gap and offset that lead to the minimal energy in the system for various combinations of *ϵ*-*A* interaction parameters. By definition, D-periodic fibrils with gap and overlap regions form when the radial gap is greater than 1.12*σ* (contact distance) whereas non-D-periodic fibrils are formed otherwise, as indicated in Figure 3b.

For the same D-mimetic molecule as investigated in our molecular dynamics simulations, in the analytical model we again find that D-periodic assemblies dominate in a wide region of the phase diagram as indicated in Figure 3c-e. Strikingly, the phase diagram that we find by this analytic model is in almost perfect agreement to the one that we gained by molecular dynamics simulations, where the region in the phase space where non-D-periodic fibrils are predicted by the analytic model coincides with the region where clusters are formed in MD simulations, as can be seen by comparing Figure 3c and Figure 1d. The value of the *radial gap* and the *offset* in the fibril determine the value of the D-periodicity. By defining the periodicity *p* again as the sum of one overlap and one gap region that appears along the backbone of the fibril, we can show that again the periodicity *p* does not vary significantly as a function of both the intermolecular interactions, as shown in Figure 3e. There, we find that the periodicity varies between *p*_max_=6.72*σ* and *p*_min_=6.03*σ* for different values of the isotropic attraction *ϵ* and the electrostatic interactions *A* with a mean value of *p*_mean_=6.37*σ*. For comparison, the MD simulations led to a mean value of *p*_mean_=5.47*σ* 0.52*σ*. The slight difference in the exact value arises as the analytic model is only one-dimensional, while the MD simulations allow for fluctuations in monomer positions in 3D.

As demonstrated in Figure 3 we have shown that our analytic model is a quick and reliable way of determining the periodicity of fibrils. Now we use this method to investigate the quantitative relationship between the charge distribution on the single molecules and the resulting periodicity of the corresponding fibrils. For that purpose we test a set of different molecule charges, as shown in

Figure 4a, and measure their periodicity for the same range of interaction parameters as shown in Figure 3e. In addition to the previously investigated D-mimetic molecule, we test three different symmetric charge distributions where both ends of the molecule possess opposite charge with various lengths of the charged regions (S1-3). We then also investigate two simple asymmetric charge distributions where the positive region typically is much greater than the negative region (A1 and A2).

**FIG. 4:**
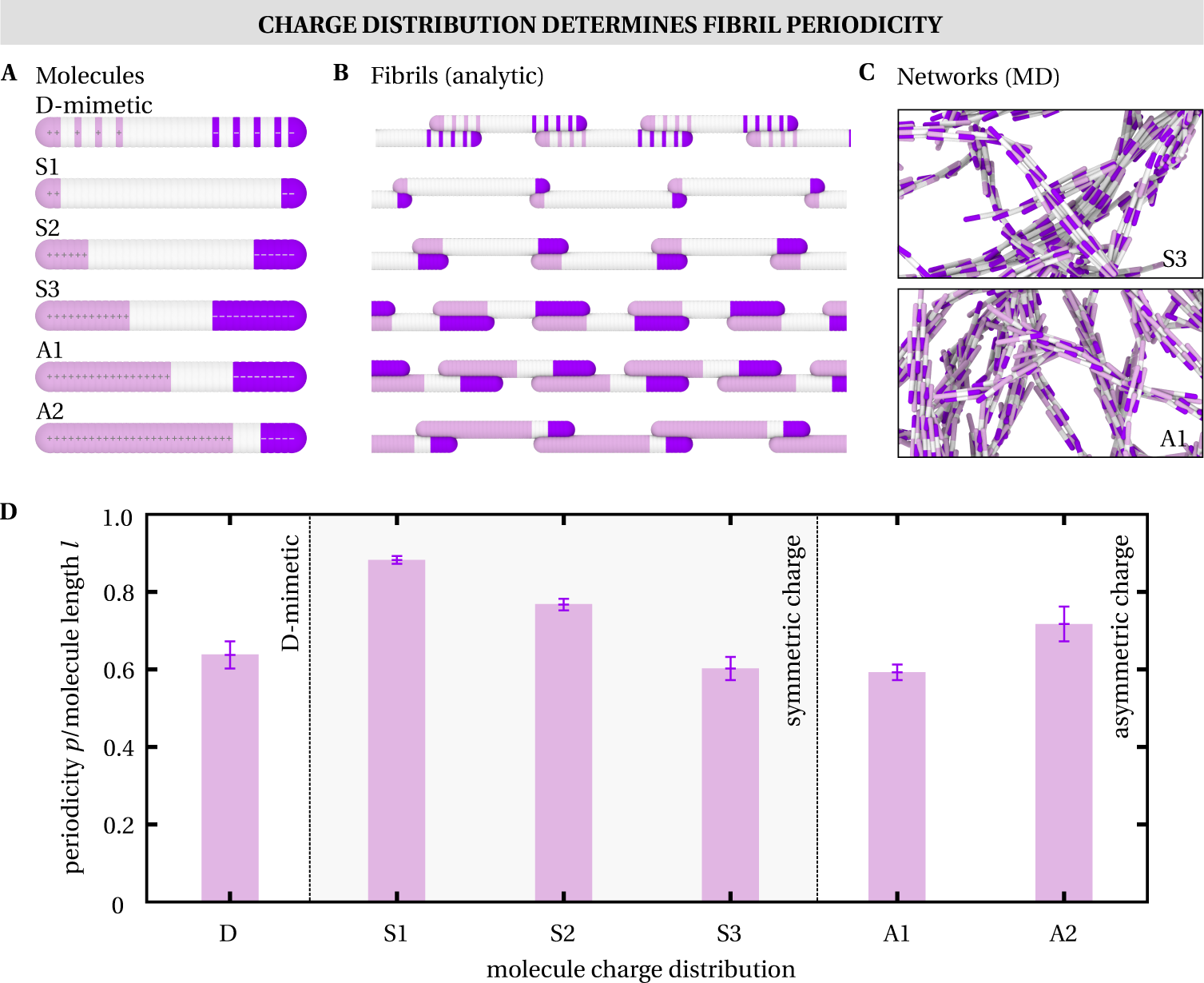
Connecting molecular design to the fibril morphologies. **a)** Different distributions of charges decorating single molecules for which we analysed the fibril periodicity. **b)** Periodic fibril configurations predicted by the analytic model for the diverse charge distributions shown in a). **c)** Corresponding MD simulations, exemplary shown for the molecules S3 and A1, show D-periodic fibril structures that are remarkably similar to the ones predicted by the analytic model as shown in b). **d)** The periodicity *p*/molecule length *l* predicted by the analytic model for the different charge distributions that are shown in a). The error bars indicate the maximum range of periodicities that are measured.

Figure 4b shows the periodic fibril configurations that are predicted by the analytic model, while Figure 4d reveals the value of the resulting periodicity where the variations indicated by the error bars are the result of the periodicity measurements for the case of different interaction parameters. Clearly, the charge distribution of individual molecules has the dominant effect on the periodicity of the fibrils, while the details of the interactions create a small variability in the value of the periodicity.

For simple symmetric molecules, such at the D-mimetic molecules and molecular designs S1-3, the value of the periodicity is effectively given by the length of the neutral region and the length of one charged region, as visible in Figure 4b. The length of the D-periodicity can be increased by making the neutral region longer. However, for just slightly more complex designs, such as asymmetric distribution of charges (A1 and A2), the situation is not as predictable any more. Here the value of the periodicity is determined not only by increasing the contact area between regions of opposite charges, but also by avoiding contacts between regions of like charges. This results in the fibril structure that cannot be easily inferred from looking at the design of a single molecule.

To confirm the results of our analytical model, we also ran MD simulations for one representative symmetric (S3) and asymmetric (A1) design. Figure 4c shows snapshots of fibrillar networks resulting from the self-assembly process in MD simulations for these molecules. The periodicity measured in simulations is again strikingly similar to the ones we predicted analytically, again confirming that the periodicity is determined solely by the pattern of charges on the molecule.

## Discussion and Conclusions

Here we developed a minimal model for simulating fibrillogenesis of collagen-mimetic molecules. The model describes a collagen-mimetic monomer as an elastic attractive rod decorated by a pattern of surface charges. When electrostatic interactions dominate over non-specific isotropic interactions, we find that fibrils of well-defined periodic structures form. The periodicity in the fibril structure is extremely robust, and resilient towards the changes in the protein concentrations, molecular flexibility, or the strength of interactions, and is determined solely by the pattern of the charged residues along the monomer backbone. Taken together, we find that the interaction parameters determine whether the fibrils will form, but once the fibrils are formed their structure is determined solely by the design of the monomer and not sensitive to the fine details of interaction parameters. We explored six different molecular designs that can be characterised in two categories based on whether the surface charge is symmetric with respect to the centre of the monomer or not. When the monomers posses symmetric charge distributions on their surface, the minimum energy configuration of two interacting monomers would be clearly the one in which the monomers are perfectly aligned in an antiparallel fashion. However, in our simulations we find that such monomers readily self-assemble into fibrils in which the monomers are aligned in a parallel fashion with a specific offset. Such structures minimise manybody interactions between many monomers and are a true emergent phenomenon.

Even this simple case demonstrates that the fibril periodicity cannot be necessarily predicted from the pair interactions of two monomers. For monomers that have more complex pattern of surface charge, such as those with asymmetric distribution, the situation is even less obvious. Interestingly we find that a simple analytical model that considers the interaction of a single monomer with periodic images of itself can be used to predict whether the fibrils will be formed and what their morphology will look like. The model predicts morphologies that are remarkably close to those obtained in molecular dynamic simulations, with slight differences arising due to local molecular fluctuations that occur in dynamic simulations. Our monomer design is indeed minimal, and reminiscent of previous synthetic collagen-mimetic molecules. We have however not explored the existence of the pattern of hydrophobic interactions or the existence of other specific interactions between two monomers that can occur in the case of native collagen molecules. It is reasonable to expect that such interactions will also play a role in determining the interactions between the monomers, hence in determining the fibril periodicity. Nevertheless, there is no reason to believe that they could not also be treated by our simple MD and analytical considerations.

Finally, it would be extremely interesting to apply this model to the case of native collagen molecules, which are much longer (∼1000 residues) and contain a more complex pattern of charged residues, and are hence beyond this proof-of-principle study. The minimal model proposed here can be also used to study the influence of the molecular design on the structures above fibrils, such as fibers and networks. We envisage that the principles put forward in this study will help guide the design of future collagen-mimetic materials and help explain the architectures of collagen fibrils found in nature, in healthy and diseased states.

## Author Contributions

A.Š. conceived the research and wrote the manuscript. A.E.H. performed and analysed computer simulations, with the input from N.G.G., C.A.B, and L.K.D. A.E.H. designed the analytical model. A.Š. supervised the project. All authors revised and approved the manuscript.

## Acknowledgments

We thank Melinda Duer, Patrick Mesquida, Lucy Colwell, Lucie Liu, and Daan Frenkel for helpful discussion. We acknowledge support from the Engineering and Physical Sciences Research Council (A.H., L.K.D. and A.Š.), BBSRC LIDo programme (N.G.G and C.B.), the Royal Society (A.Š), and the UK Materials and Molecular Modelling Hub for computational resources, which is partially funded by EPSRC (EP/P020194/1).

